# Behavioral context determines network state and variability dynamics in monkey motor cortex

**DOI:** 10.1101/233684

**Authors:** Alexa Riehle, Thomas Brochier, Martin Paul Nawrot, Sonja Grün

## Abstract

Variability of spiking activity is ubiquitous throughout the brain but little is known about its contextual dependence. Trial-to-trial spike count variability, estimated by the Fano Factor (FF), and within-trial spike time irregularity, quantified by the local coefficient of variation (CV2), reflect variability on long and short time scales, respectively. We co-analyzed FF and CV2 in monkey motor cortex comparing two behavioral contexts, movement preparation (*wait*) and execution (*movement*). We find that FF significantly decreases from *wait* to *movement*, while CV2 increases. The more regular firing (low CV2) during *wait* is related to an increased power of local field potential beta oscillations and phase locking of spikes to these oscillations. In renewal processes, a widely used model for spiking activity under stationary input conditions, both measures are related as FF≈CV^2^. This expectation was met during *movement*, but not during *wait* where FF≫CV2^2^. We conclude that during movement preparation, ongoing brain processes result in changing network states and thus in high trial-to-trial variability (high FF). During movement execution, the network is recruited for performing the stereotyped motor task, resulting in reliable single neuron output. We discuss our results in the light of recent computational models that generate non-stationary network conditions.

## Introduction

A number of *in vivo* studies have demonstrated a high variability of single neuron spiking activity in the neocortex. Here we address the question whether and how cortical spiking statistics within a trial and across trials may depend on the behavioral context aiming at an improved understanding of the underlying nature and sources of neuronal variability. In our experimental analyses we co-analyze two types of spiking statistics that reflect variability on separate time scales (Nawrot et al. 2008; Murray et al. 2014). *Spike time irregularity* (Perkel et al. 1967; Softky and Koch 1993) refers to the random appearance of a sequence of spikes. This finds quantitative expression in the dispersion of inter-spike intervals (ISI) within a trial, statistically captured by the *coefficient of variation* (CV) of ISIs. It represents variability on a relatively short time scale, in the range of tens to a few hundreds of milliseconds, determined by the typical duration of ISIs (Nawrot et al. 2008; Softky and Koch 1993; Holt et al. 1996). *Trial-by-trial spike count variability* measures the variation in the number of spikes across repeating experimental trials of the same behavioral condition. It can statistically be quantified by the *Fano factor* (FF; Shadlen and Newsome 1998; Nawrot et al. 2008) defined as the ratio between the variance and the mean of spike counts measured across trials within an observation window of predefined length. It represents variability on a rather long time scale in the range of seconds determined by the typical separation of trials that belong to an identical behavioral condition. For both measures, high CV and FF values indicate irregular ISIs and high spike count variability, respectively.

Spike time irregularity is much lower in the sensory periphery (Werner and Mountcastle 1963) or motor periphery (Calvin and Stevens 1968; Clamman 1969; Prut and Perlmutter 2003; Barry et al. 2007; Duclos et al. 2008) than in cortical primary sensory and motor areas. Within cortical areas, the CV decreases systematically from visual to higher-order sensorimotor areas (Maimon and Assad 2009; Shinomoto et al. 2009; Mochizuki et al. 2016). Likewise, the FF differs with stages of sensory processing, being lowest in the periphery and highest in the cerebral cortex (Kara et al. 2000).

Other studies have explored the modulations of spike time irregularity and spike count variability in relation to behavior. Davies et al. (2006) showed that in motor cortex spike time irregularity increases during the most demanding epoch of a precision grip task. Spike count variability, on the other hand, systematically decreases during a behavioral task being lowest during movement execution (Churchland MM et al. 2006, 2010; Rickert et al. 2009), perceptual processing (Mitchell et al. 2009; Churchland MM et al. 2010; Abolafia et al. 2013; Ponce-Alvarez et al. 2013; Mazzucato et al. 2015), attention (Mitchell et al. 2007; but see McAdams and Maunsell 1999), or in relation to decision processes (Churchland AK et al. 2011). In addition, Rickert et al. (2009) showed that during movement preparation and execution the FF in motor cortex depends on the amount of prior information about the requested movement. However, these studies suffer important limitations. First, they do not systematically assess how changes in spiking variability relate to changes in firing rate. Second, these studies analyzed spike time irregularity and spike count variability separately although *in vitro* studies suggest that these two measures may be directly related (Nawrot et al. 2008).

Here we take advantage of a very large database of single neuron recordings from the motor cortex of three macaque monkeys to decipher the relationship between firing rate, spike time irregularity and spike count variability. In particular, we demonstrate how this relationship changes in different behavioral contexts, specifically during movement preparation and execution.

## Materials and Methods

### Behavioral task

Three adult macaque monkeys (*Macaca mulatta*), two females (monkey L and E), weighing 4.5 and 7kg, and one male (monkey N), weighing 7 kg, were used in the experiment. All animal procedures were approved by the local ethical committee (authorization A1/10/12) and conformed to the European and French government regulations.

Details of the task and recording procedures were described in Riehle et al. (2013) and Milekovic et al. (2015). Monkeys were trained to perform an instructed delay reach-to-grasp task to obtain a food reward (apple sauce), using the left hand. They sat in a custom-made primate chair in front of the experimental apparatus with the non-working (right) arm loosely restrained in a semi-flexed position. The unrestrained working hand rested on a switch positioned at waist-level, 5 cm lateral to the midline. The target object was a stainless steel, rectangular parallelepiped (40×16×10 mm) attached to the anterior end of a low-friction horizontal shuttle and rotated at a 45° angle from the vertical axis. It was located 13 cm away from the switch at 14 cm height. The object had to be grasped and pulled with the working hand using one of two different grip types: a precision grip (PG) or a side grip (SG). The object weight could be set to one of two different values (100 or 200g) by means of an electromagnet inside the apparatus. Thus, the force required to pull the object was either low (LF) or high (HF). Changes in object weight occurred between trials and were undetectable by the monkey. The apparatus provided a continuous measure of the grip and pulling (load) forces by means of force sensitive resistances (FSR). In addition, a hall-effect sensor measured the horizontal displacement of the object over a maximal distance of 15 mm.

A square of 4 red light-emitting diodes (LEDs) with 1 additional yellow LED in its center was used to display the instruction cues. The LEDs were inserted in the apparatus just above the target object. Illumination of the two left or right red LEDs instructed the monkey to perform a SG or a PG, respectively. Illumination of the two bottom or top LEDs instructed the monkey that pulling the object required a LF or HF, respectively.

The task was programmed and controlled using LabView (National Instruments Corporation, Austin, TX, USA). The trial sequence was as follows. The monkey had to close the switch with the hand to self-initiate a trial (trial start, TS). After 400 ms, the central yellow LED was illuminated for another 400 ms (warning signal, WS), followed by the preparatory cue (Cue), illuminated for 300 ms, which instructed the monkey about the grip (PG or SG) required to perform the trial. Cue extinction was followed by a 1 s preparatory delay. At the end of this delay, the GO signal provided the remaining information about the force and also served as the imperative signal asking the monkey to release the switch (switch release; SR) and to reach and grasp the object. Following object grasp, the monkey had to pull the object towards him/her into a narrow position window (4-14mm) and to hold it there for 500 ms to obtain the reward (Rew). In case of grip error, the trial was aborted and all four LEDs were flashed as a negative feed-back. The reaction time (RT) was defined as the time between the GO signal and SR, and the movement time (MT) as the time between SR and grip force onset as detected by the FSR by using a fixed threshold. The monkey was required to keep RT and MT below 700 ms, for monkeys L and N, and 1000ms for monkey E to be rewarded. Five to 10 sessions of about 10 to 15 minutes each were recorded per day, up to five recording days per week. During each session, the four trial types (SG-LF, SG-HF, PG-LF, PG-HF) were presented at random with equal probability. Monkeys usually achieved a total of 100 to 140 successful trials per session. Only one session per recording day was selected for analysis to strongly reduce the probability to analyze the same neurons twice (see more details below).

### Surgery

When the monkey was fully trained in the task and obtained 85% correct trials, a 100-electrode Utah array (Blackrock Microsystems, Salt Lake City, UT, USA) was surgically implanted in the motor cortex contralateral to the working hand. The array had an arrangement of 10x10 Iridium Oxide electrodes, each of them 1.5mm long, with an inter-electrode distance of 400 μm. The surgery was performed under deep general anesthesia using full aseptic procedures. Anesthesia was induced with 10mg/kg i.m. ketamine and maintained with 2-2.5 % isoflurane in 40:60 O_2_-air. To prevent cortical swelling, 2 ml/kg of mannitol i.v. was slowly injected over a period of 10 minutes. A 20x20 mm craniotomy was performed over the motor cortex and the dura was incised and reflected. The array was positioned on the cortical surface 2-3mm anterior to the central sulcus at the level of the spur of the arcuate sulcus (see Fig. 1A). The array was inserted using a pneumatic inserter (Array Inserter, Blackrock Microsystems) and covered with a sheet of an artificial non-absorbable dura (Preclude, Gore-tex). The live dura was sutured back and covered with a piece of an artificial absorbable dura (Seamdura, Codman). The bone flap was put back at its original position and attached to the skull by means of a 4×40 mm strip of titanium (Bioplate, Codman). The array connector was fixed to the skull on the opposite side with titanium bone screws (Codman). The skin was sutured back over the bone flap and around the connector. The monkey received a full course of antibiotics and analgesics before returning to the home cage.

**Figure 1:**
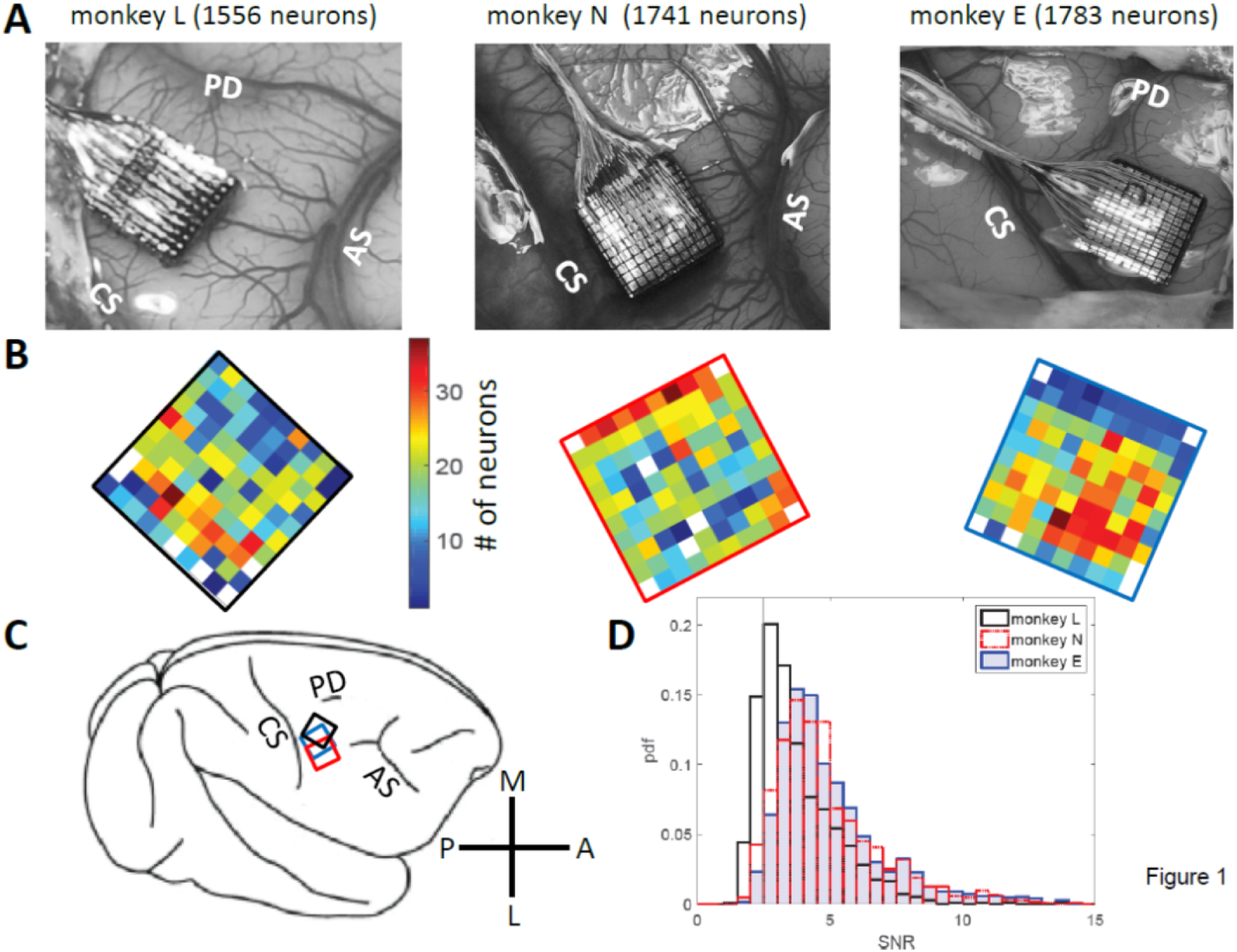
Data set and electrode array locations. **A:** Locations on the cortical surface of the Utah arrays implanted in monkey L (left), N (middle), and E (right), all in the right hemisphere (see C). Pictures were taken during surgery. CS: central sulcus; AS: arcuate sulcus; PD: precentral dimple. **B:** Numbers of selected neurons on each electrode, SNR>2.5. Same color code (# of neurons) for both monkeys (monkey L at the left with a black frame, monkey N at the middle with a red frame, and monkey E at the right with a blue frame). **C**: A schematic drawing of the right hemisphere including the placement of the arrays for each monkey using the same colors as for the frames shown in B. M: medial; A: anterior; L: lateral; P: posterior. **D**: Distributions of SNR values for all neurons for all three monkeys. Vertical line corresponds to the selected SNR threshold value (2.5).

### Recordings

Neuronal data were recorded using the 128-channel Cerebus acquisition system (NSP, Blackrock Microsystems). The signal from each active electrode (96 out of the 100 electrodes were connected) was preprocessed by a head stage (monkey L: CerePort plug to Samtec adaptor, monkeys N and E: Patient cable, Blackrock Microsystems) with unity gain and then amplified with a gain of 5000 using the Front End Amplifier (Blackrock Microsystems). The raw signal was obtained with 30kHz time resolution in a range of 0.3Hz to 7.5kHz. From this raw signal, two filter settings allowed us to obtain two different signals by using filters in two different frequency bands, the local field potential (LFP, low-pass filter at 250Hz) and spiking activity (high-pass filter at 250 Hz). The LFPs were sampled at 1 kHz and saved on disk. On each channel, the experimenter set online a threshold for detection and extraction of potential spikes. All waveforms crossing the threshold were sampled at 30 kHz and snippets of 1.6 ms duration for monkey L and 1.3 ms for monkey N and E were saved for offline spike sorting. All behavioral data such as stimuli, switch release, force traces for thumb and index fingers and object displacement were also fed into the Cerebus, sampled at 1 kHz and stored for offline analysis.

### Data set, selection criteria and analysis windows

Single neurons were sorted offline from the online extracted waveforms by using the Plexon Offline Spike Sorter (version 3.3.3, Plexon Inc., Dallas, TX, USA). Spike clusters which were separated significantly from each other and with less than 1 *%* of inter-spike intervals (ISIs) of durations of ≤ 2 ms were considered as single neurons. For further selection, we calculated from the spike shapes of each neuron the signal-to-noise ratio (SNR), defined as the amplitude, i.e. the mean of the trough-to-peak voltage, divided by twice the standard deviation of the entire signal (Hatsopoulos et al. 2004). For this study we selected only single neurons with SNR values > 2.5 to guarantee good sorting quality. The distributions of the SNR values of all neurons are shown in Fig. 1D. For further analysis, spike times were down-sampled to 1 kHz.

For data analysis, we selected representative sessions from the entire recording period. The criteria for selection were (i) a large number of recorded neurons (64 – 110 neurons in monkey L, 92 – 167 neurons in monkey N, 121 – 165 neurons in monkey E), and (ii) a continuous behavior throughout the session without interruption between trials. From monkey L, 21 sessions were selected, recorded from Oct. 5, 2010 until April 15, 2011, including 1929 single neurons in total. Of those, 1556 neurons remained with SNR>2.5 and were included in the analysis. From monkey N, 13 sessions were selected, recorded from June until Nov. 2014, including 1826 single neurons, of those 1741 neurons with SNR>2.5 were selected for the subsequent analysis. And finally, from monkey E, 13 sessions were selecetd, recorded during Dec. 2016 and Jan. 2017, including 1828 neurons, of those 1783 neurons with SNR>2.5 were selected for the subsequent analysis. A small fraction of these neurons were likely to be the same across different sessions (Dickey et al. 2009). For the number of neurons selected on each electrode across all selected sessions, see Fig. 1B. Note that for the study presented here simultaneity of the recorded spike data is not relevant, but it was relevant to achieve a large sample. The sessions selected in monkey L were: l101005-002, l101006-002, l101007-001, l101008-003, l101013-002, l101014-002, l101015-001, l1011108-001, l101110-003, l101111-002, l101126-002, l101202-001, l101209-001, l101216-002, l101220-002, l110208-001, l110209-001, l110404-001, l110408-002, l110411-001, l110415-002; in monkey N: i140613-001, i140616-001, i140617-001, i140627-001, i140701-001, i140702-001, i140703-001; i140704-001, i140718-001, i140721-002, i140725-002, i140917-002, i141117-001, and in monkey E: e161209-001, e161212-002, e161213-001, e161214-001, e161215-001, e161216-001, e161219-002, e161220-001, e161222-002, e170105-002, e170106-001, e170109-001, e170110-004.

### Computing variability measures

To analyze variability measures as a function of the behavioral context, we selected two discrete 500ms epochs. To study neuronal activity during *wait*, data were aligned to the GO signal and the first 500ms of the preparatory delay were selected, starting at Cue offset (horizontal thick black bars in Fig. 2; left panels). We selected this specific window, because firing rate in most neurons was most stable during this period. For the analysis of neuronal activity during *movement*, data were aligned to switch release (SR in the right panels, i.e. aligned to movement onset; in the left panels this event was indicated as avSR, meaning average SR times for data aligned to GO). In monkeys L and N a window from 150ms before to 350ms after SR was selected, whereas in monkey E a window from SR to 500ms after SR was selected. These windows covered most of the movement period represented by the peak of average firing rate after GO (green horizontal bars in Fig. 2, right panels). Note that monkey E had much longer RTs than the two other monkeys, see inset in Fig. S1.

**Figure 2:**
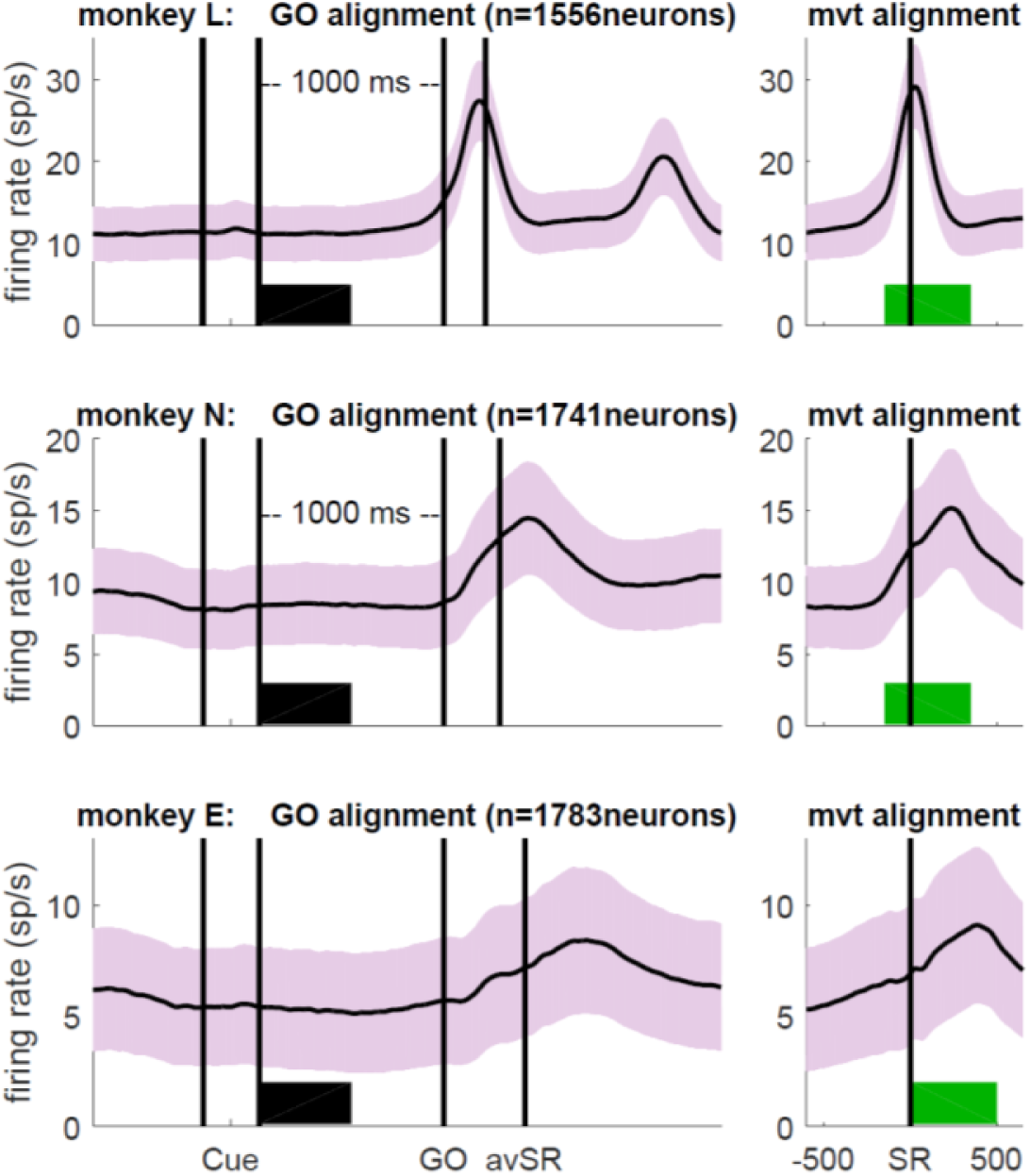
Firing rates averaged across all selected neurons. (SNR>2.5) during correct trials for monkey L (top), monkey N (middle) and monkey E (bottom), obtained during the trial type SG-HF. Light colored envelopes indicate SEM. Data were aligned to the GO signal (left) and movement onset (SR; right). The selected analysis windows of 500ms duration are indicated by a thick bar in black (*wait*) and green (*movement*). Cue: preparatory signal providing prior information about the grip type; GO: Go signal; SR: switch release (i.e., movement onset); avSR: average SR for data aligned to the GO signal. For firing rates recorded during each trial type as well as reaction times, see Fig. S1.

The CV of ISIs is defined as

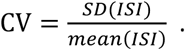

This measure is meaningful only for a constant firing rate and largely overestimates the irregularity of spiking activity if firing rate changes in time (see Ponce-Alvarez et al. 2010). To deal with non-stationary firing rates we used instead a local measure, the CV2 that was introduced by Holt et al. (1996). It is based on neighboring ISIs (*m*-values). Individual *m*-values were computed for any two consecutive ISIs as

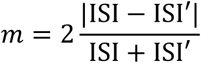

where *m* takes values between 0 (no variability) and 2 (maximum variability). Each *m*-value is associated with the time of occurrence of the second of the three considered spikes. For each analysis epoch (*wait* or *movement*), the CV2 is obtained by averaging across all *m*-values extracted in all trials that fall within the selected analysis window

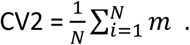

As a consequence, the 1^st^ or the 3^rd^ spike belonging to the two consecutive ISIs determining a respective *m*-value may be outside of the window. Reliability of the estimates requires a minimum of 20 *m*-values per window across all trials (see Ponce-Alvarez et al. 2010), thus segments with a lower number of *m*-values were not considered. We repeated the central analysis of our study using a different local measure of irregularity, the LV (Shinomoto et al. 2005), yielding the same qualitative results (see also Ponce-Alvarez et al. 2010).

The spike count variability was measured by the Fano factor (FF; Softky and Koch 1993; Stevens and Zador 1998) defined as

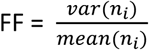

i. e. the ratio between the variance and the mean of the spike counts *n* across trials *i* within the analysis epoch (*wait* or *movement*). In order to minimize the estimation bias for the FF due to a finite window size we did not consider samples with less than 5 spikes/s per epoch. For processes that are more regular than Poisson, the bias then becomes negligible (Nawrot et al. 2008; Nawrot 2010).

By theoretical argument, for any given stochastic point process the irregularity of ISIs (CV) and the variability of the spike count (FF) are related. Specifically for the model class of renewal processes, where ISIs are independent and equally distributed, the prediction is CV^2^ ≈ FF (Cox and Lewis 1966; Perkel et al. 1967; Shadlen and Newsome 1998; Chacron et al. 2001; Nawrot et al. 2008; Ditlevsen and Lansky 2011; Farkhooi et al. 2011). In both measures, a Poisson process corresponds to a value of 1 and lower/higher values to lower/higher variable spiking activity. The prediction CV^2^ ≈ FF seems to be fulfilled *in vitro* (Stevens and Zador 1998; Nawrot et al. 2008), but not *in vivo* as observed in the awake behaving monkey (Nawrot 2010). In the latter stationarity across trials is not given, such that conclusions on the renewalty of the data cannot be drawn.

### Phase-locking of spikes to LFP beta oscillations

Phase-locking of spikes to LFP beta oscillations was analyzed in extended windows of 1000ms duration. For *wait*, the window was selected from 500ms before until 500ms after cue offset, and for the *movement* epoch a window was selected from 400ms before until 600ms after movement onset. In order to exclude trivial signal correlations between the occurrence of spikes and the phase of the LFP oscillation induced by volume conductance effects (Katzner et al. 2009), we averaged trial by trial for each electrode channel the LFP signals recorded from all directly neighboring channels (between 4 and 9 channels as a function of its location on the array). From this averaged LFP signal, the maximum frequency in the beta range was determined between 10 and 45 Hz from the power? Context-related variability in spiking activity spectrum during the enlarged *wait* window. This frequency was then used to filter the LFP signals throughout the entire length of the behavioral trial with a zero-phase bandpass filter (Matlab *filtfilt*, max. frequency ± 5Hz, Butterworth, 4 pole). From this filtered signal, we calculated the instantaneous phase of the LFP obtained via a Hilbert transformation (Matlab *hilbert*; see Fig. 6B). We then related for each neuron recorded on that electrode its spike times separately during each of the two analysis windows to the beta phase of the LFP (Matlab *angle*). In order to obtain the significance of a possible phase-locking of the spikes to the beta oscillation (p<0.05), we used the Rayleigh test for non-uniformity of circular data in each window (CircStat Toolbox for Matlab, *circ_rtest*; Berens 2009). For each analysis window, only neurons were taken into account having more than 30 spikes, making that the number of neurons varied between the two windows.

## Results

We explored in motor cortex the relationship between spike time irregularity and trial-by-trial spike count variability and analyzed the dependency of these two measures to changes in firing rate. For the spike time irregularity, we used instead of the classical coefficient of variation of ISIs a local measure, the CV2, that was introduced by Holt et al. (1996) in order to deal with non-stationary firing rates. For the spike count variability, we determined the Fano factor, FF (see Materials and Methods). Three monkeys (L, N and E) were trained in a delayed reach-to-grasp task (Riehle et al. 2013; Milekovic et al. 2015) in which the animal had to grasp, pull and hold an object using either a side grip (SG) or a precision grip (PG) and employing either a low force (LF) or high force (HF), thus resulting in a total of 4 trial types (SG-LF, SG-HF, PG-LF, PG-HF). These trial types were presented to the animal in a pseudo-random fashion from trial to trial with equal probability in each recording session. We explored potential changes of the two variability measures, CV2 and FF, as a function of the behavioral context by selecting two contextually different epochs during the performance of the task. As a first epoch we chose a 500ms-window directly after cue offset during movement preparation, subsequently called *wait*. During this epoch the monkey was not allowed to move but was requested to prepare the precued movement based on the information provided by the cue. The second epoch was chosen during active movement execution, subsequently called *movement*. In each monkey, the 500ms-window was roughly centered around the moment of the average peak discharge in the firing rate histogram of all neurons across all trials (see Fig. 2). In monkeys L and N the 500ms-window was chosen from 150ms before to 350ms after SR, whereas in monkey E who reacted more slowly to the GO signal, the 500ms-window was chosen starting with SR (see Fig. 2). We only considered single neuron activity (i) from trials of correct behavior, and (ii) with a high signal-to-noise ratio (SNR>2.5) of the spike shapes (see Materials and Methods). Thus, the analysis in monkey L was performed on 1556 neurons (21 sessions), in monkey N on 1741 neurons (13 sessions), and in monkey E on 1783 neurons (13 sessions) (Fig. 1B, for the number of neurons on the electrode arrays). Fig. 2 shows the trial-average firing rates of all selected neurons for each of the three monkeys during trials of one out of the four trial types (SG-HF).

For all our analyses, we aligned the neuronal activity to the GO signal (Fig. 2, left) and to movement onset (i.e. SR; Fig. 2, right) in the *wait* and the *movement* epoch, respectively. In the left panels of Fig. 2 the two first vertical lines indicate the onset and offset of the preparatory cue, presented for 300ms, that provided prior information about the grip type (here SG). The third vertical line indicates the occurrence of the GO signal that provided the missing information about the force to pull the object (here HF). The GO signal also requested the execution of the reach-to-grasp movement. Since the reaction time (RT) is variable across trials, SR times were averaged over all correct trials within each session, and indicated in Fig. 2 (left panels) by a vertical line after GO as avSR (average switch release). The performance speed of the monkeys was different. Their average RTs across all trial types and sessions were 170ms, 257ms and 413ms for monkey L, N and E, respectively. The second phasic rate increase visible in monkey L was due to her movements back to the center key, and is visible here only because her RTs were fast. For RTs obtained in each trial type for each monkey, see inset in Fig. S1.

We performed all analyses separately for each of the four trial types. However, since the results did not differ between them, we only show in the following the results for SG-HF, as in Fig. 2. The results obtained during the other trial types are shown in the Supplementary Information.

### Irregularity (CV2) and trial-by-trial variability (FF) are modulated with the behavioral context

In a first step of our analysis we determined whether and how neuronal variability modulated with the behavioral context. For each single neuron we computed the spike time irregularity (CV2) and the spike count variability across trials (FF) separately in the two selected epochs *wait* and *movement*. For each epoch we considered only neurons that fulfilled our selection criteria for variability analysis, i.e. a minimum trial-averaged firing rate of 5 spikes/s and a sufficient number of ISIs to reliably compute CV2 and FF within the 500ms window (see Materials and Methods). All averages are expressed by the median. We find that both types of variability are modulated with the behavioral context. The CV2 is lower during *wait* than during *movement,* whereas the FF shows the opposite behavior (Fig. 3). These contextual modulations are statistically highly significant (Wilcoxon ranksum test, p<10^−4^) and are consistent across the three monkeys and across all four trial types (Table S1). Additionally, the firing rate significantly increases from *wait* to *movement* (Fig. 3 and Table S1).

**Figure 3:**
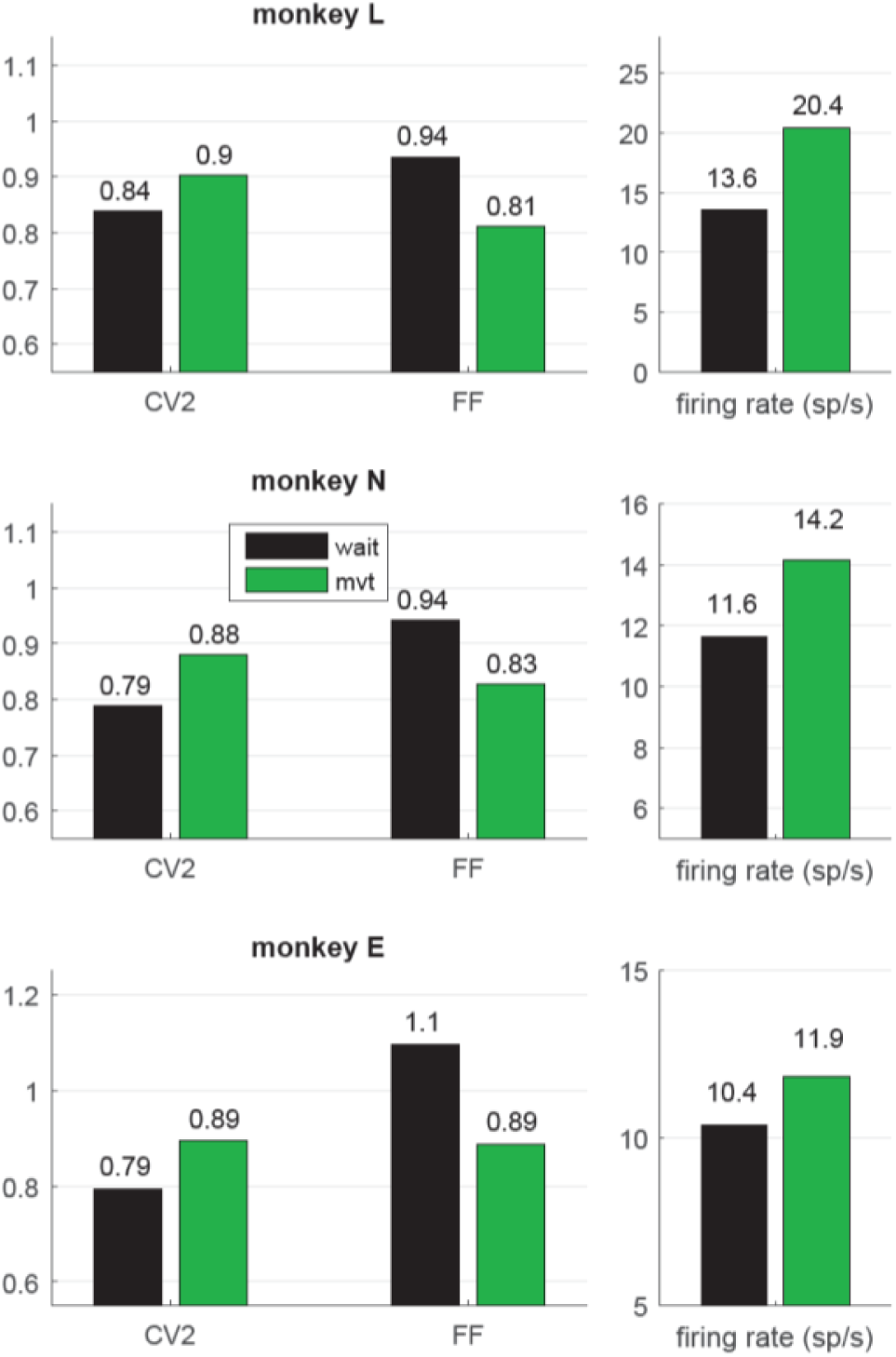
Context-dependent modulations of CV2, FF and firing rate during *wait* and *movement*. All values are indicated as medians and are significantly different (Wilcoxon ranksum test; p<10^−4^) between *wait* (black) and *movement* (green) for all three monkeys. For the number of neurons, see Table 1. Data in this figure were obtained during the trial type SG-HF, values determined in all trial types are provided in Table S1.

**Table 1:** Selection of data sets, number of neurons (SG-HF)

|  | monkey L<br>*wait* | monkey L<br>*mvt* | monkey N<br>*wait* | monkey N<br>*mvt* | monkey E<br>*wait* | monkey E<br>*mvt* |
| --- | --- | --- | --- | --- | --- | --- |
| all recorded neurons / selected with SNR>2.5 | 1929 / 1556 |  | 1826 / 1741 |  | 1828 / 1783 |  |
| selected neurons with > 5sp/s in either epoch (Fig. 3) | 981 | 1313 | 1000 | 1215 | 583 | 821 |
| of those selected with respect to mean activity (act) (Fig. 4):<br>$act_{mvt} > act_{wait} \mid act_{wait} > act_{mvt}$ | 758\|223 | 1127\|186 | 622\|378 | 953\|262 | 336\|247 | 676\|145 |
| selected neurons with > 5sp/s and > 20 *m*-values for CV2 in both epochs (Figs. 5 and S2) | 944 |  | 884 |  | 481 |  |
| selected neurons for phase-locking analysis with >30 spikes per analysis window (Fig. 6) | 1245 | 1408 | 1048 | 1167 | 848 | 1063 |

In order to explore the behavior of the individual neurons we analyzed the spiking activity of all neurons and selected the ones that fulfilled the selection criteria in both epochs. This drastically reduced the number of neurons (944, 884 and 481) in monkeys L, N and E, respectively (see Table 1). For those we inspected for each single neuron and each of the measures (CV2, FF and firing rate) its value observed during *wait* against that obtained during *movement* (Fig. 4A-C). The obtained scatter diagrams show for each measure that in each monkey the majority of individual neurons indeed changed their behavior from *wait* to *movement*, consistent with the results gained from averaging across neurons (shown in Fig. 3). In all three monkeys 67 % of the neurons increased the CV2, whereas about 60-70 % of the neurons decreased the FF, and for a vast majority of neurons (70-80 %) the firing rate increased from *wait* to *movement*. This is summarized in Fig. 4D by using the contrast value

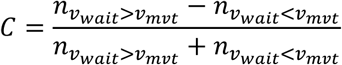

where the two expressions correspond to the number of neurons with a higher value during *wait n*_*v*_*wait*>*v*_*mvt*___ and the other with a higher value during *movement n*_*v*_*wait*>*v*_*mvt*___. This contrast measure signifies with a positive outcome a higher portion of higher values during *wait*, and *vice versa*, with a negative outcome a higher portion of higher values during *movement*.

**Figure 4:**
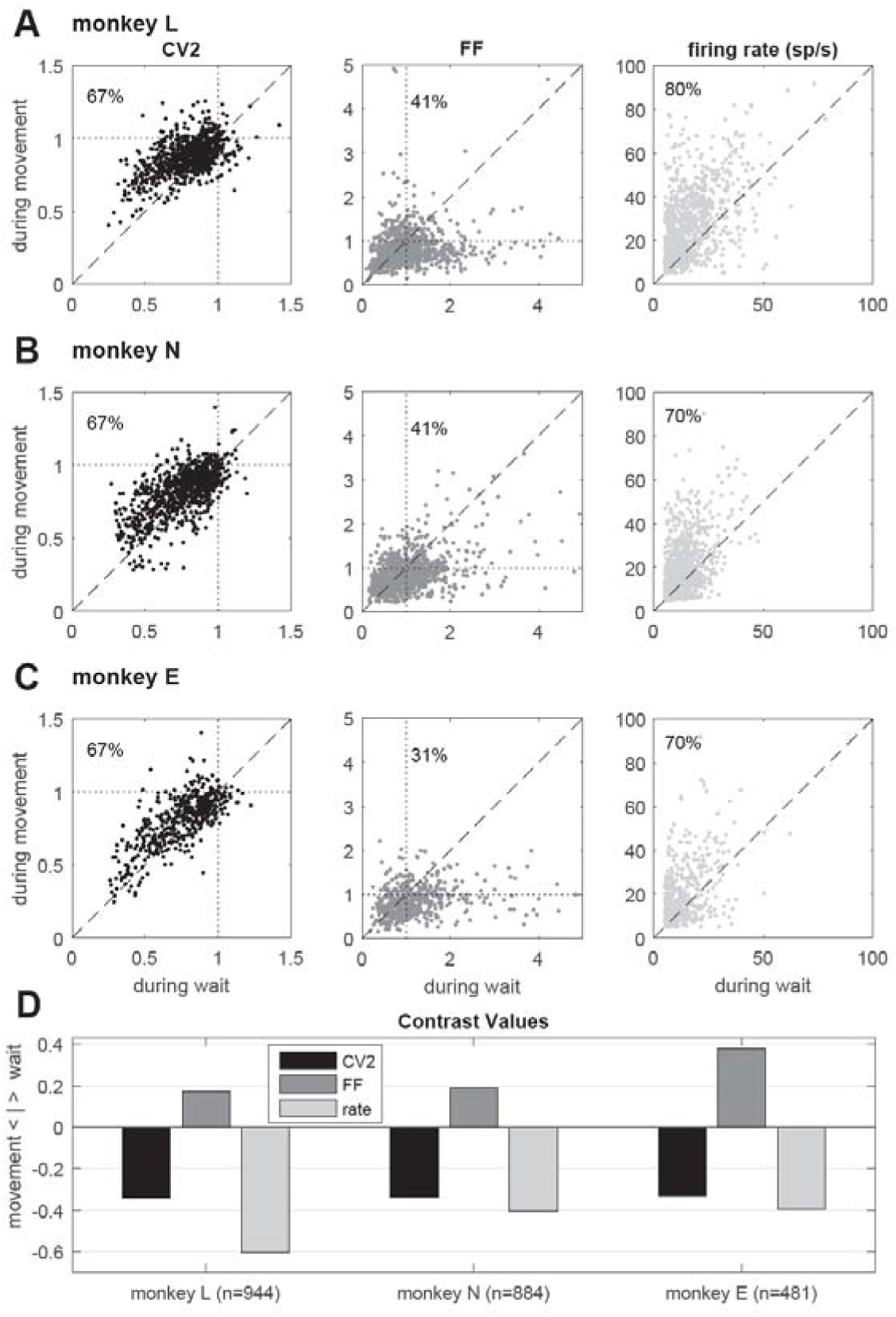
Single neuron comparison of variability measures and firing rate. **A-C:** Scatterplots of CV2 (in black), FF (in dark gray) and firing rate (in light gray) obtained during *wait* (x-axis) and *movement* (y-axis) for monkey L (n=944), N (n=884) and E (n=481) for neurons fulfilling the selection criteria in both epochs. The number in the upper left corner indicates the percentage of neurons whose values are higher during *movement* than during *wait*. **D**: Contrast values for each feature. Data in this figure were obtained during the trial type SG-HF, for results in the other trial types see Fig. S2.

We found this result to be consistent for all trial types (see Fig. S2).

As evident from Fig. 2, the time-resolved firing rate averaged across all neurons shows a clear increase during *movement* (see also Fig. 3). However, in a minority of neurons (20-30%) the firing rate decreased during *movement* as compared to the *wait* epoch (Fig. 4A-C, right panels). We therefore explored whether the modulation of variability depends on these differential firing rate modulations. For that purpose we split the neurons into two subpopulations with respect to their firing rate profile during the two epochs. One subpopulation contained the majority of neurons with a firing rate that was higher during *movement* than during *wait* (80%, 70% and 70% in monkeys L, N and E, respectively; see Table 1), whereas the other subpopulation contained the remaining neurons with a higher firing rate during *wait* than during *movement*. We again found that the CV2 generally increases and FF generally decreases from *wait* to *movement* irrespective of the firing rates of the individual neurons (see Fig. 5 for SG-HF and Table S2 for data from all monkeys and trial types).

**Figure 5:**
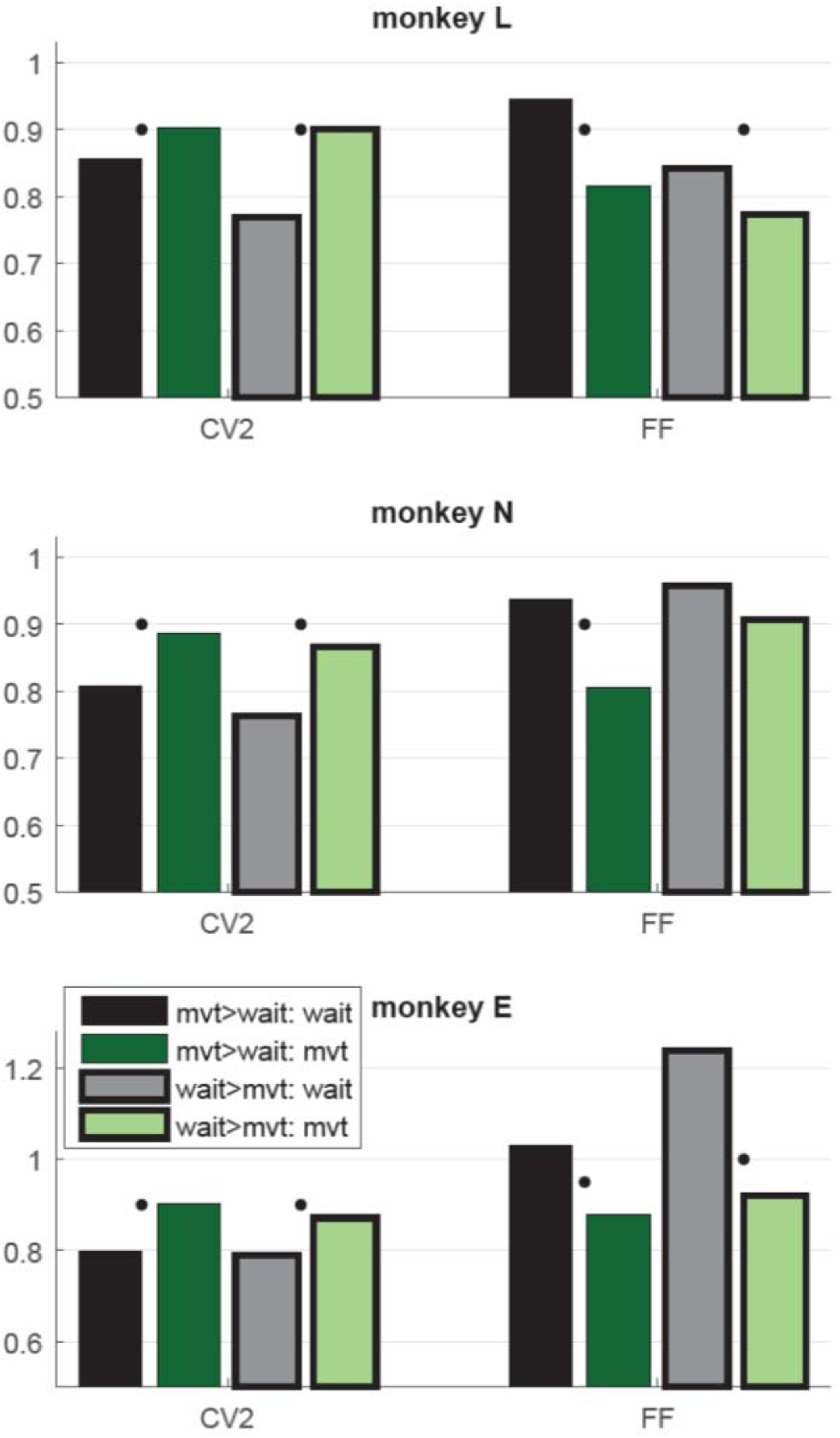
Context-dependent modulations of CV2 and FF during *wait* and *movement* as a function of the differential firing rate during the two task epochs. Data from trial type SG-HF. Variability measures obtained in two subpopulations of neurons, those with a higher firing rate during *movement* than during *wait* (mvt>wait; no surrounding, two left bars) and vice versa (wait>mvt; black surrounding, two right bars). For the numbers of neurons in each subpopulation, see Table 1. Medians of CV2 and FF during *wait* and *movement* for each subpopulation. For each measure and each subpopulation, the significant differences between *wait* and *movement* are indicated by a black dot (Wilcoxon ranksum test; p<0.05). Data in this figure were obtained during the trial type SG-HF, for data obtained in all four trial types, see Table S2.

### Increased spike time irregularity coincides with reduced phase-locking of spikes to the LFP

Next, we explored why the CV2 is lower during *wait* than during *movement* (as shown in Fig. 3). A possible reason for the more regular firing during *wait* than during *movement* may lie in the tendency of spikes to lock to LFP oscillations (Denker et al. 2007, 2011). The power of motor cortical LFP beta oscillations (15-35 Hz) strongly increases during an instructed delay, but is lowest during movement execution (Sanes and Donoghue 1993; Pfurtscheller et al. 1996; Kilavik et al. 2012; for a review, see Kilavik et al. 2013). We therefore hypothesized that a majority of neurons exhibit phase locking of their spikes to the beta oscillations during *wait* but not during *movement* where beta oscillations are vanishing. This would imply, as a consequence, that the spiking activity is more regular during *wait* than during *movement*.

Therefore, we analyzed LFP oscillatory activity in the beta range and related the spike times of each single neuron to the oscillation phase of the LFP signal. For each recording channel, we first averaged trial-by-trial the LFP signals recorded from all direct neighboring channels to exclude trivial signal correlations between the occurrence of spikes and the phase of the LFP oscillation induced by volume conductance effects (Katzner et al. 2009). Fig. 6A shows a spectrogram obtained from such a LFP signal in a single session in monkey N, averaged across all trials of trial type SG-HF. A clear beta oscillation occurred during cue presentation and the subsequent delay period. The beta oscillation ceased during movement execution and only re-occurred around the reward (Rew) at the end of the trial. We next determined the spike occurrences of each neuron recorded on each electrode with respect to the beta phase of the LFP oscillation averaged across all neighboring electrodes (see Materials and Methods). Fig. 6B shows during 5 single trials the raw (averaged) LFP signals in gray, the beta-filtered signal (main beta frequency 18Hz ± 5Hz) in black, and the instantaneous oscillation phase in cyan. The spike times of one single neuron recorded on the center electrode are indicated by red dots.

**Figure 6.**
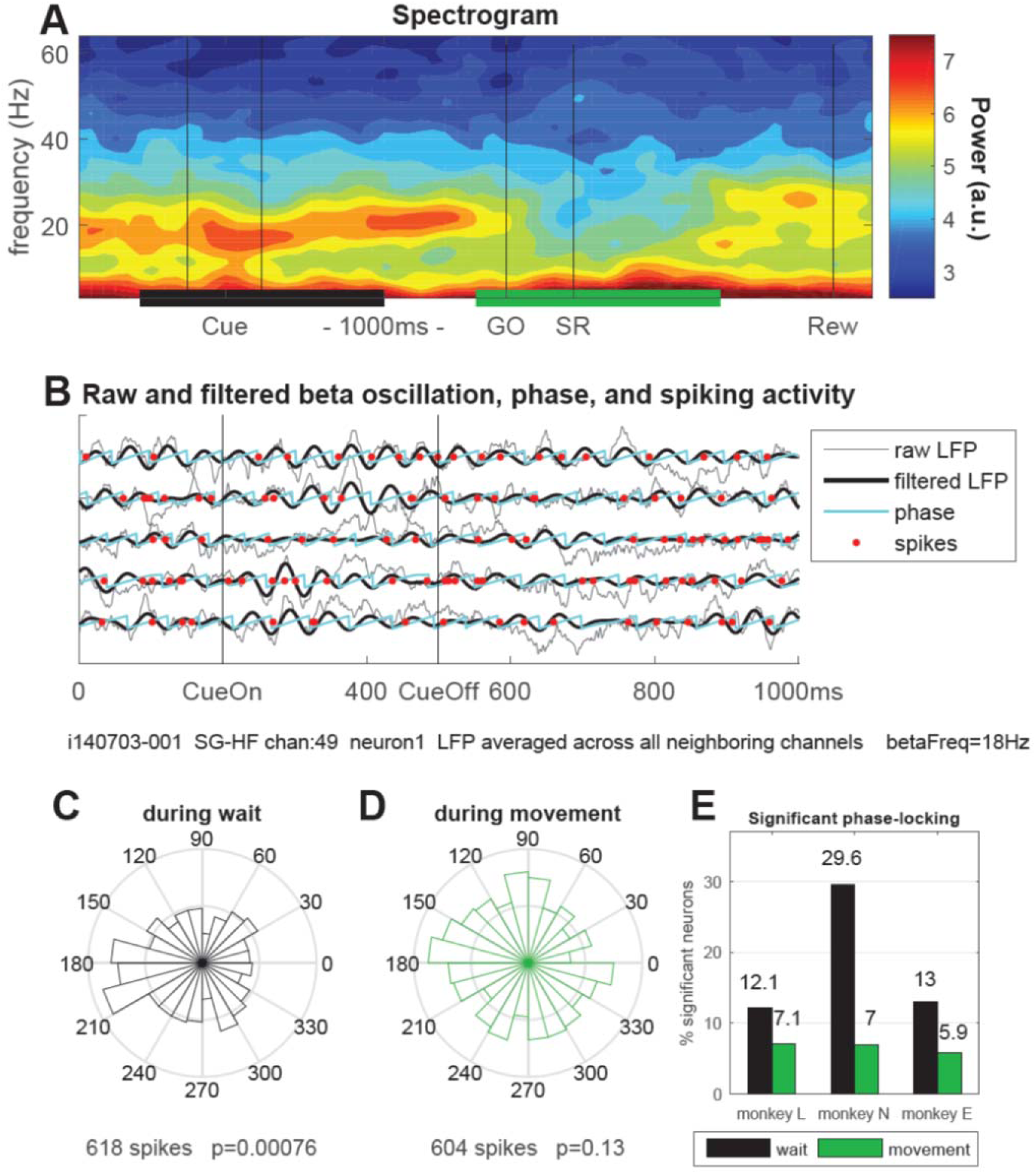
Phase-locking of spiking activity to LFP beta oscillations. **A:** Spectrogram of the LFP signal averaged trial-by-trial across all neighboring electrodes around the reference electrode (channel 49) and then across all behavioral trials (n=36) of a single trial type (here SG-HF) during one recording session in monkey N (i140703-001). The power is indicated as a color code in arbitrary units. The black and green horizontal bars at the bottom indicate the selected windows for calculating the phase-locking of spikes to the LFP oscillation phase. **B:** The LFP and spiking data recorded on electrode 49 during 5 selected trials are shown, cut around cue presentation from 500ms before until 500ms after cue offset (CueOff). The duration of the cue was 300 ms. The raw (averaged) LFP signals are shown in gray. The signals in black were band-pass filtered around the main beta oscillation frequency (here 18Hz ±5Hz; see Methods). The oscillation phases obtained by Hilbert transformation are plotted in cyan. The spike times of one single neuron (neuron 1) recorded on channel 49 are indicated by red dots. **C** and **D**: Circular representation of the phase relationships of the spiking activity of a single neuron obtained during *wait* (C) and during *movement* (D), same neuron as the one presented in B by red dots, but recorded during all SG-HF trials. The number of spikes during the respective analysis window and the p-value of the outcome of the Rayleigh test are shown on the x-axis of the figures. **E**: Percentages of statistically significant (p<0.05) phase-locked neurons during *wait* (black) and *movement* (green) in the trial type SG-HF in monkeys L, N and E, respectively. For the number of selected neurons see Fig. S3 (upper panel). The results of the same analysis performed during all other trial types are shown in Fig. S3.

We then determined for each neuron the phase values at its spike times in each of the two selected epochs, *wait* and *movement*, and calculated their phase distributions in circular space (see Fig. 6C and D). To capture potential phase locking of the spike times we evaluated the statistical significance (p<0.05) of the non-uniformity of each distribution using the Rayleigh test (see Fig. 6C, D, for the example data). Repeating this analysis for all neurons showed indeed a higher percentage of significantly phase-locked neurons during *wait* (black) than during *movement* (green), in all three monkeys (Fig. 6E). Fig. S3 shows that the relative fractions of significantly phase-locked neurons during *wait* and *movement* which were almost the same in all trial types and for each monkey. Thus, phase-locking of spikes is prominent during *wait* but almost at chance level during *movement*.

### Context-modulated relationship of spike time irregularity and cross-trial spike count variability

For the final step of our analysis we considered predictions from stochastic point process theory (see Materials and Methods). For a given point process model, spiking irregularity (CV) and count variance (FF) are related in a unique manner. Under certain conditions, spike time irregularity and spike count variability are related as FF ≈ CV^2^ (Stevens and Zador 1998; Shadlen and Newsome 1998; Nawrot et al. 2007, 2008; Ditlevsen and Lansky 2011). This assumes stationary conditions and is true for all renewal processes, where ISIs are independent and equally distributed, and which are widely used as models for spiking activity (Perkel et al. 1967; Tuckwell 1988; Chacron et al. 2001; Nawrot 2010). The statistical equality of FF ≈ CV^2^ was confirmed in *in vitro* experiments that used stationary noise current injection to stimulate single cortical neurons (Stevens and Zador 1998; Nawrot et al. 2003a,b; Nawrot et al. 2008).

We tested this relationship for the spiking data of our *in vivo* recordings and plotted the squared local measure, CV2^2^, against the FF. Fig. 7 shows these scatter diagrams separately for each monkey and behavioral epoch. We found that the expectation for the hypothesis (see Materials and Methods) was met in the *movement* epoch (Fig. 7, right panels, in green), where all data points are close to the diagonal (on average FF/CV2^2^ = 0.96, 1.11 and 1.14 for monkeys L, N and E, respectively). The scattering around the diagonal can easily be explained by the variance of estimation, because we have only a limited number of trials per neuron (Nawrot 2010). During *wait* (Fig. 7, left panels, in black), however, there was a strong and systematic deviation from the diagonal, where FF was by far larger than CV2^2^ (on average FF/CV2^2^ = 1.43, 1.6 and 1.94 for monkeys L, N and E, respectively). In the *wait* epoch we found only a minority of single neurons (20%, 17% and 12% in monkeys L, N and E, respectively) for which FF was smaller than CV2^2^, i.e. FF/CV2^2^ < 1, whereas during *movement* context this held true for approximately half of the neurons (55%, 40% and 38% in monkeys L, N and E, respectively). Table S3 shows that both the percentages of neurons with a smaller FF than CV2^2^ and the average ratios (median) of FF/CV2^2^ during *wait* and *movement* were very similar in all four trial types and all three monkeys. Our results indicate that during *wait* the theoretic prediction for renewal processes is clearly violated in a systematic manner whereas during *movement* the neurons match the renewal expectation FF ≈ CV2.

**Figure 7:**
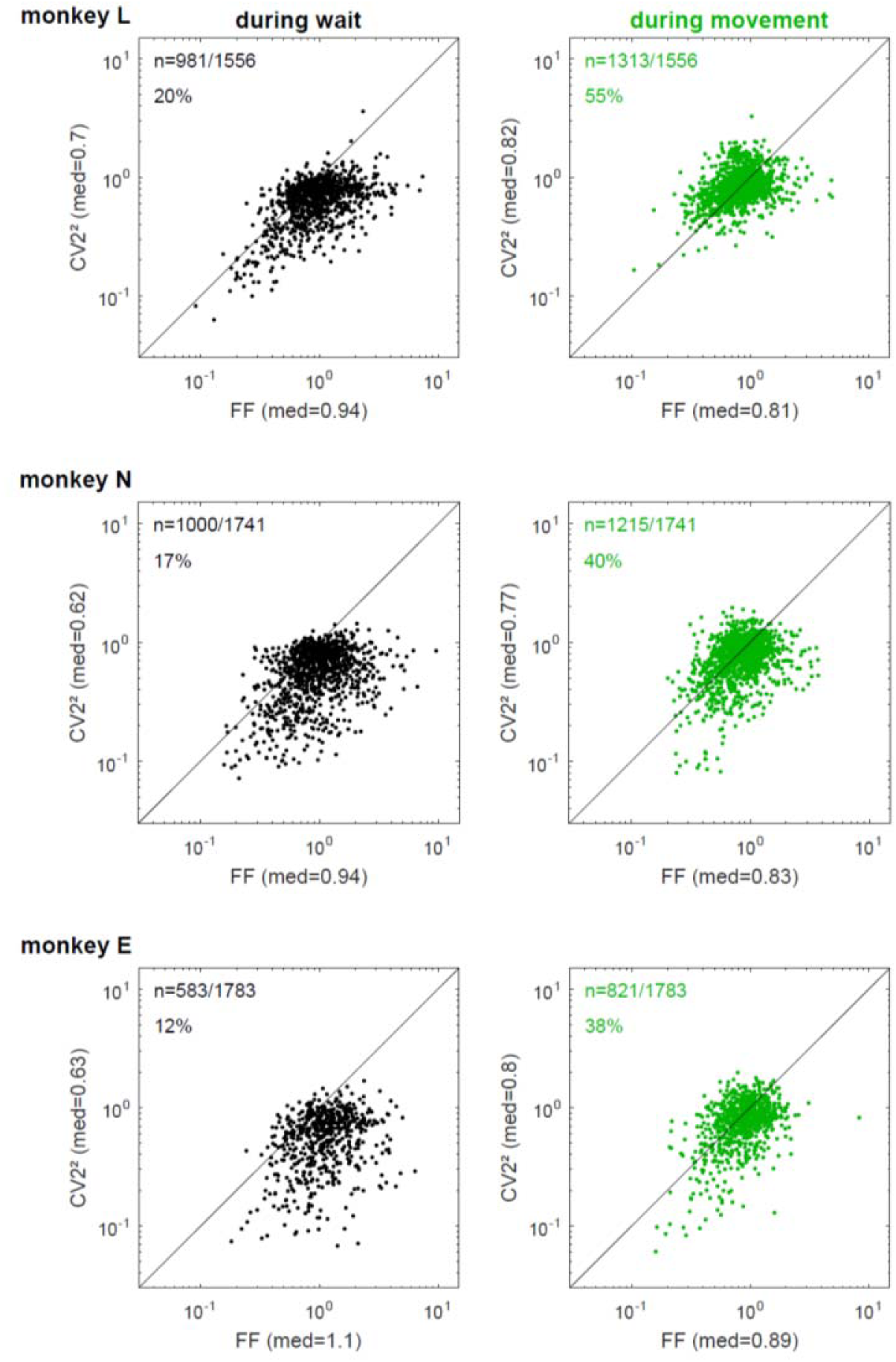
The relationship of spike time irregularity and spike count variability depends on the behavioral context. Log-log representation of scatter diagrams of the CV2^2^ vs FF during *wait* (black, left) and movement (green, right) for each monkey in SG-HF The medians of FF and CV22 are indicated on each axis. In the left upper corner of each plot, the number of selected neurons and the percentage of neurons is indicated whose ratio FF/CV2^2^ was smaller than 1. These percentages as well as the the ratio FF/CV2^2^ during all trial types and for all monkeys are shown in Table S3.

## Discussion

The present study provides new insights into the variability of single neuron spiking activity and their dependence on the behavioral context. In line with previous studies on trial-by-trial variability in motor cortex (Nawrot et al. 2003a; Churchland MM et al. 2006, 2010; Rickert et al. 2009), our results show that the FF decreases strongly from movement preparation (*wait*) to execution (*movement*). The same task-related dynamics of across-trial count variability was confirmed for different cortical areas in a large meta-study (Churchland MM et al. 2010). In addition, we here report two new important findings. First, as opposed to the FF, the CV2 is on average significantly higher during *movement* than during *wait* (Fig. 3), irrespective of the firing rate (Fig. 5). Second, we found that the relation FF ≈ CV2^2^ for each neuron is fulfilled during *movement*, but not during *wait* where the FF was considerably larger than the CV2^2^ (Fig. 7). Both results, detailed for one trial type (SG-HF) in the Results section, are consistent in all other trial types (see Supplementary Information, Tables S1 and S4).

### Context-dependent modulation of spike time irregularity

Our data show that the CV2 increases from *wait* to *movement.* This is in agreement with Davies et al. (2006) who found during a precision-grip task a considerable fraction of pyramidal tract neurons exhibiting more regular firing during hold than during movement execution. Similarly, Compte et al. (2003) showed in prefrontal neurons a modulation of CV2 as a function of task requirements, with a lower CV2 during eye fixation than during a more demanding mnemonic delay. Our results cannot be simply explained as an artefact of the firing rate modulation during *movement* as shown in ground truth simulation studies where similar rate modulations did not impact the CV2 (Ponce-Alvarez et al. 2010; Hamaguchi et al. 2011).

During *wait* our data show a low spike time irregularity as expressed by a low CV2. These results point to a strong link between the low CV2 and the high LFP beta power. Indeed, we find that during *wait* the spiking activity of a larger percentage of neurons is significantly phase-locked to LFP beta oscillations than during *movement* (see Fig. 6E and S4). This is in agreement with the findings that high power LFP beta oscillations (15-35Hz) were observed during *wait*, whereas during *movement* beta oscillations were lowest (see Fig. 6A; Sanes and Donoghue 1993; Pfurtscheller et al. 1996; Kilavik et al. 2012, 2013). Murthy and Fetz 1992, 1996) described that many motor cortical neurons tend to fire in phase with high amplitude LFP oscillations, but they did not provide any indication of regular firing in absence of LFP oscillations. Furthermore, beta power was also increased during active hold in a precision-grip task compared to movement execution (Baker et al. 1997; Davies et al. 2006). Irregularity is shown to be lower in all periods of elevated beta oscillation, as described in the cited literature, than during periods with weak or no beta oscillations.

### Modulation of FF vs. CV2 indicates context-dependent change of the cortical network state

During *movement,* our data robustly obey the renewal prediction FF ≈ CV^2^. Theoretical considerations and *in vitro* experiments indicate that under stationary input conditions cortical neurons are close to the renewal prediction. This was confirmed in acute rat cortical slice preparations when single neurons were stimulated with stationary noise currents (Nowak et al. 1997; Stevens and Zador 1998; Nawrot et al. 2003a, b, 2007, 2008). Increasing the input noise, for instance, through balancing excitation with inhibition, increases overall variability of the output spike trains albeit with a fixed relation of FF ≈ CV^2^. Furthermore, *in vivo* intracellular recordings in anesthetized rat somatosensory cortex showed that FF is equal or even slightly smaller than the CV^2^ (FF ≤ CV^2^) under stationary conditions (Nawrot et al. 2007). In visual cortex of behaving monkeys, Brostek et al. (2013) showed that spiking statistics of MSTd neurons, both during the presentation of moving visual stimuli and during ocular following, are in good agreement with the renewal prediction. Unfortunately the authors did not test the renewal prediction before stimulus presentation and/or eye movements, which would have been better suited for comparison with our results during *wait*.

During *wait* we found a strong deviation from the theoretic prediction for renewal processes with FF ≫ CV2^2^. This confirms preliminary results on a different and smaller data set of monkey motor cortical neurons analyzed during a period of movement preparation (Nawrot 2010). There are two possible interpretations of this result (see Materials and Methods). The spiking activity of individual neurons might deviate from the renewal statistics during *wait*. This interpretation is unsatisfying because the very same neurons show a renewal-like spiking statistics during *movement*. In addition, spiking statistics close to renewalty has also been demonstrated for cortical neurons *in vitro* that were stimulated with injection of stationary noise currents (see above). We clearly favor the alternative interpretation which is a violation of the stationarity assumption implying varying input conditions that change from trial to trial. We hypothesize that the large trial-by-trial variability *in vivo* during *wait* reflects a trial-to-trial change of the network state implying a trial-to-trial change of the single neuron input level. Indeed, mimicking non-stationary input conditions *in vitro* by noise current injections where the input noise level was mildly varied across trials resulted in a strongly increased FF ≫ CV^2^ (Nawrot et al. 2003c).

What could be the cause for the postulated change in network state across trials? Recent theoretical models provide phenomenological and mechanistic explanations for the observed modulation of the FF that support our interpretation of non-stationary conditions across trials. A simple phenomenological point-process model combined a renewal process with slow and moderate across-trial variation of the point process intensity (i.e. the single neuron firing rate) to account for ongoing network activity that is not task-related (Arieli et al. 1996). This resulted in a spontaneous spiking statistics where FF ≫ CV^2^. Adding a stereotyped task-related activation component that mimics activation during movement resulted in a reduction of FF such that FF ≈ CV^2^ (Nawrot 2010). Recent studies of large-scale clustered spiking neural network models suggested an interesting mechanistic explanation for the task-related FF reduction. Neuron clusters formed by highly interlinked groups of excitatory neurons result in a network activity that cycles through different attractor states (Deco and Hugues 2012; Litwin-Kumar and Doiron 2012; Mazzucato et al. 2015). Combining excitatory and inhibitory neuron clusters further increases the robustness of the observed attractor dynamics. In spontaneous conditions this cluster cycling effectively leads to a permanent change in the network state. In neural network simulations with spiking neurons that mimic an experimental trial design this cluster dynamics results in a high single neuron spike count variability (high FF) before stimulation. However, when a particular cluster is stimulated, the network activity becomes focused on this cluster, while activity in non-stimulated clusters is suppressed through global inhibition. Trial-by-trial stimulation of the same cluster results in a strongly reduced trial-to-trial variability during stimulation in all neurons (Deco and Hugues 2012; Litwin-Kumar and Doiron 2012; Mazzucato et al. 2015). This model could reproduce the task-related FF dynamics observed *in vivo* (Churchland MM et al. 2010). However, these studies currently lack a detailed analysis of spiking irregularity that would allow for a thorough comparison with physiology.

An alternative model explanation is based on the self-inhibiting cellular mechanism of spike frequency adaptation (SFA) that is ubiquitous in spiking neurons including cortical neurons (Lundstrom et al. 2008). If model neurons in a cortical network with balanced excitation and inhibition are equipped with a SFA current, stimulation of an excitatory population leads to a strong stimulus-locked transitory reduction of FF that qualitatively and quantitatively fits the physiological *in vivo* observations (Farkhooi et al. 2013). Both suggested mechanisms - attractor dynamics at the network level and SFA at the cellular level - are likely to act in concert in order to produce the complex state and context-dependent FF dynamics observed in experimental data (Rickert et al. 2009; Churchland MM et al. 2010; Churchland AK et al. 2011).

### Conclusions

Motor cortex is particularly well suited to explore how the dynamics of spiking activity is modulated by the behavioral context. We conclude with the following interpretation: During *wait,* regularity expressed by a low CV2 is related to the increased power of LFP beta oscillations. Across-trial variability expressed by the FF, however, is high because in each trial the local network is in a different state implying a different amount of synaptic input and spike output of single neurons for each trial. This non-stationarity across trials was attributed to ongoing brain activity (Arieli et al. 1996). Indeed, motor cortical activity is modulated long before movement execution by processes such as visuomotor transformation (Di Pellegrino and Wise 1993; Riehle et al. 1997; Zhang et al. 1997) or movement preparation (Riehle 2005; Confais et al. 2012). These specific trial-by-trial modulations are under the influence of upstream activity from sensory and associative areas and may be caused by attractor cycling in clustered cortical network models (Deco and Hugues 2012; Litwin-Kumar and Doiron 2012; Mazzucato et al. 2015). Thus we observe that FF is high while CV2^2^ is low during *wait* simply because we sample spiking of individual neurons across different network states. Thus the assumption of stationarity, which is implicit to the renewal assumption, is violated. Single neuron spiking might still represent renewal spiking albeit with a slightly different process rate in each trial (Nawrot 2010). During *movement*, the local motor cortical network is recruited for performing the motor task. Single neurons receive strong task-related and stereotyped input. In effect, in each trial the neuronal population activity reliably occupies the same optimal subspace related to the desired movement (Churchland MM et al. 2006; Shenoy et al. 2013) and single neuron output shows a reduced trial-by-trial variability. The CV2, however, is enhanced due to the lack of phase locking, such that we find an agreement of FF and CV2.

## Acknowledgments

The authors thank Moritz Helias, Farzad Farkhooi and Thomas Rost for valuable discussions, Yaoyao Hao, Margaux Duret and Justine Facchini for their help during recordings, Ivan Balansard for surgical expertise, the animal house team for animal care, and Xavier Degiovanni and Joel Baurberg for technical assistance. This work was supported by the Collaborative Research Agreements CNRS-RIKEN and the CNRS-FZ Jülich to AR and SG; Agence National de la Recherche (ANR-GRASP) to TB and AR; BrainScaleS (EU Grant 269912) to SG and AR; International Associate Laboratory (LIA) “Vision-for-Action” to AR, TB, SG; Helmholtz Portfolio “Supercomputing and Modeling for the Human Brain (SMHB)” to SG and AR; EU Grants 720270 (HBP) and 604102 (HBP) to SG, and German Israeli Foundation grant “Multiple time scales of signals and noise in the motor hierarchy” (GIF grant I-1224-396.13/2012) to MN.

---

**Figure S1.**
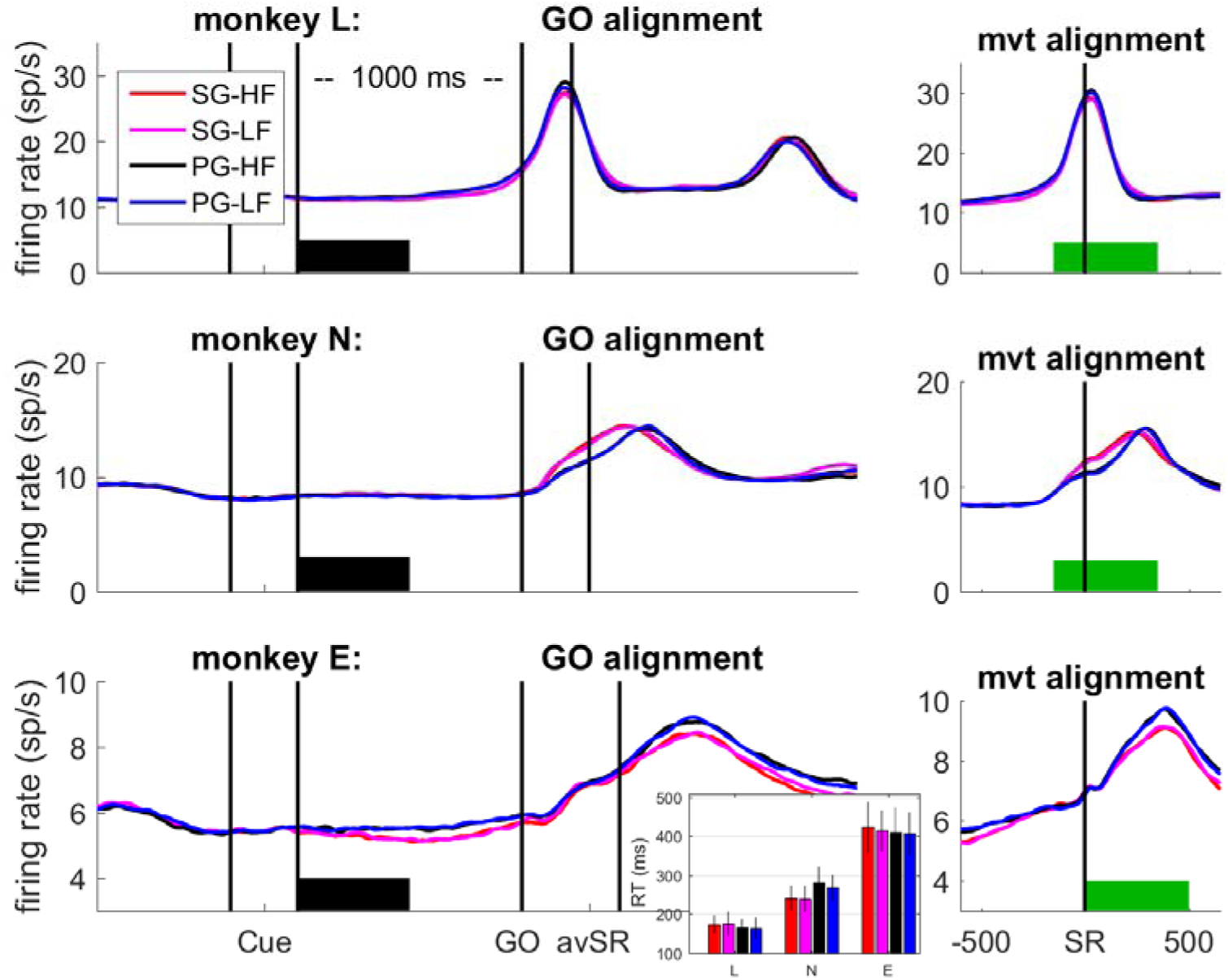
**Task-related firing rates averaged across all selected neurons** SNR>2.5) during correct trials for monkey L (top), monkey N (middle) and monkey E (bottom) during all trial types (see Fig. 2 for SG-HF). Data were aligned to the GO signal (left) and movement onset (switch realease, SR; right). The selected analysis windows of 500ms duration are indicated by a thick bar in black (*wait*) and green (*movement*). SG: side grip; PG: precision grip; HF: high force; LF: low force; Cue: preparatory cue; GO: Go signal; SR: switch release (movement onset). Number of neurons monkey L: n=1556, monkey N: n=1741, and monkey E: n=1783. It can clearly be seen that there were hardly no differences in firing rates between conditions for the entire data sets. The inset in the lower panel shows the average reaction times (RT), i.e. the time between GO and SR, for each monkey and all four trial types.

**Table S1:** **CV2, FF, and firing rate during *wait* and *movement* during all trial types**. All averages are medians. For each feature, differences between *wait* and *movement* are highly significant (Wilcoxon ranksum test; p<10^−4^). Firing rate is in spikes/second.

| monkey | trial type | CV2 wait | CV2 mvt | FF wait | FF mvt | rate wait | rate mvt |
| --- | --- | --- | --- | --- | --- | --- | --- |
| L | SG-HF | 0.84 | 0.9 | 0.94 | 0.81 | 13.6 | 20.4 |
| L | SG-LF | 0.84 | 0.9 | 0.94 | 0.81 | 13.5 | 20.9 |
| L | PG-HF | 0.83 | 0.9 | 0.88 | 0.79 | 13.9 | 20.8 |
| L | PG-LF | 0.83 | 0.9 | 0.86 | 0.8 | 14 | 20.8 |
| N | SG-HF | 0.79 | 0.88 | 0.94 | 0.83 | 11.6 | 14.2 |
| N | SG-LF | 0.79 | 0.88 | 0.95 | 0.85 | 11.6 | 14.2 |
| N | PG-HF | 0.8 | 0.88 | 0.98 | 0.86 | 11.4 | 13.2 |
| N | PG-LF | 0.8 | 0.88 | 0.98 | 0.86 | 11.4 | 13.4 |
| E | SG-HF | 0.77 | 0.88 | 1 | 0.87 | 10.5 | 11.6 |
| E | SG-LF | 0.78 | 0.88 | 1 | 0.88 | 10.6 | 11.5 |
| E | PG-HF | 0.78 | 0.88 | 1.1 | 0.92 | 10.4 | 11.9 |
| E | PG-LF | 0.8 | 0.9 | 1.2 | 0.95 | 10.4 | 11.8 |

**Figure S2:**
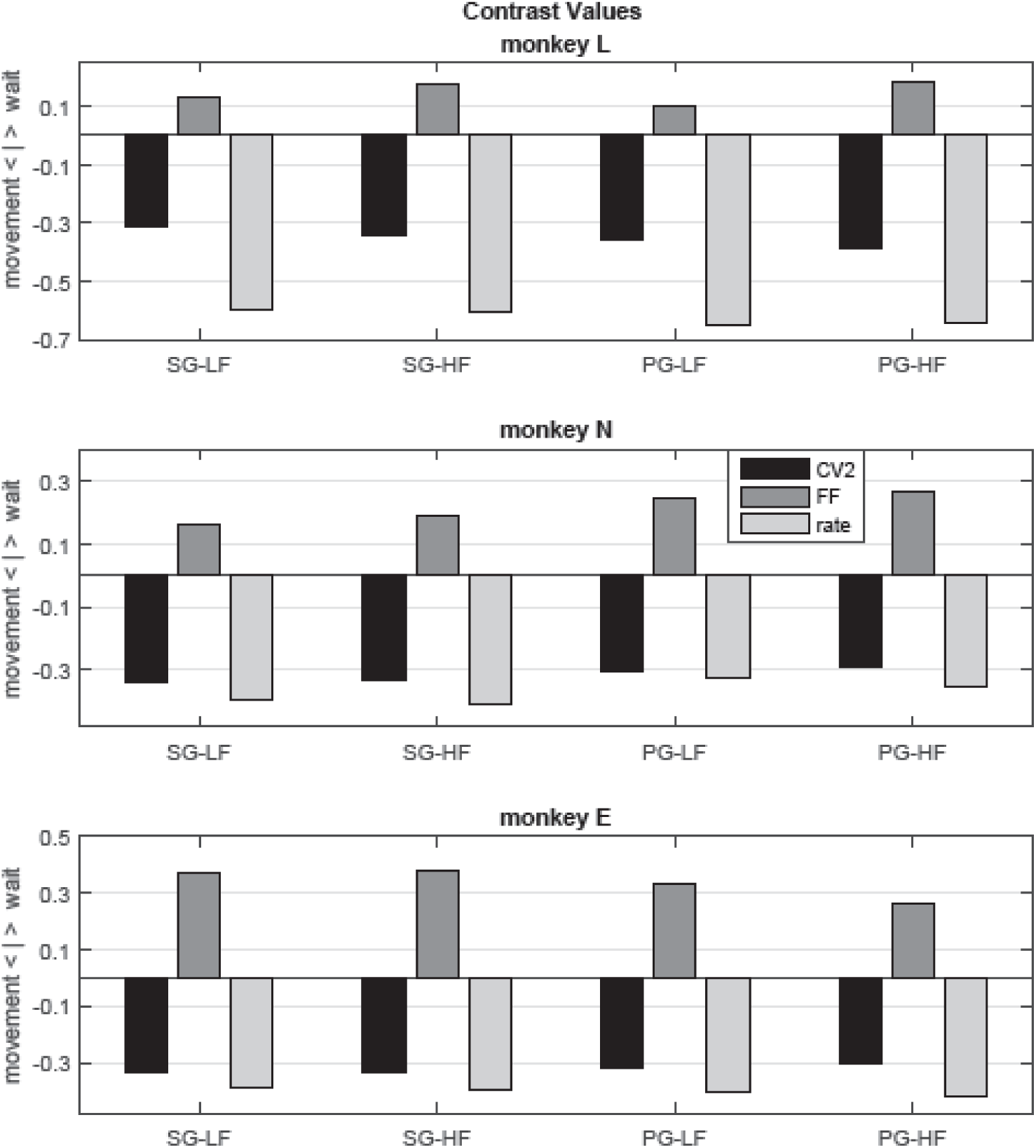
Contrast values for each measure (CV2, FF and firing rate) between *wait* and *movement*. Data obtained for each trial type and each monkey.

**Table S2:** **CV2, FF and firing rate as a function of the behavioral context and firing rate profile (see also Fig. 5).** We defined two types of firing rate profiles: the firing rate profile 1 had a firing rate during *movement* being larger than that during *wait* [act(mvt)>act(wait)], and *vice versa* the firing rate profile 2 had a firing during *wait* being larger than that during *movement* [act(wait)>act(mvt)]. Statistical significance of the differences each of the three measures between *wait* and *movement* is indicated by * in the row between the wait and movement columns (p<0.05; Wilcoxon ranksum test). All CV2 differences between *wait* and *movement* were statistically significant and CV2 was always lower during *wait* than during *movement.* For rate profile 1, all FF differences were statistically significant and the FF was always higher during *wait* than during *movement*. However for rate profile 2, differences were not always significant, most likely due to the small number of neurons in these subpopulations (see Table 1). Note, that rate differences were by definition statistically significant, because they served as selection criterion.

| monkey | trial type | CV2<br>wait | sig | CV2<br>mvt | FF<br>wait | sig | FF<br>mvt | rate<br>wait | sig | rate<br>mvt |
| --- | --- | --- | --- | --- | --- | --- | --- | --- | --- | --- |
| L profile 1 | SG-HF | 0.86 | * | 0.9 | 0.94 | * | 0.82 | 12.9 | * | 22.3 |
| L profile 1 | SG-LF | 0.85 | * | 0.9 | 0.95 | * | 0.81 | 12.9 | * | 22.8 |
| L profile 1 | PG-HF | 0.85 | * | 0.91 | 0.9 | * | 0.8 | 13.2 | * | 22.1 |
| L profile 1 | PG-LF | 0.84 | * | 0.91 | 0.89 | * | 0.81 | 13.7 | * | 23 |
| L profile 2 | SG-HF | 0.77 | * | 0.9 | 0.84 | * | 0.77 | 15.9 | * | 12.3 |
| L profile 2 | SG-LF | 0.77 | * | 0.89 | 0.89 | * | 0.79 | 15.4 | * | 11.9 |
| L profile 2 | PG-HF | 0.73 | * | 0.9 | 0.75 | * | 0.73 | 16.5 | * | 11.9 |
| L profile 2 | PG-LF | 0.75 | * | 0.89 | 0.76 |  | 0.75 | 14.8 | * | 11.5 |
| N profile 1 | SG-HF | 0.81 | * | 0.89 | 0.94 | * | 0.8 | 11.0 | * | 16.2 |
| N profile 1 | SG-LF | 0.81 | * | 0.88 | 0.95 | * | 0.83 | 10.9 | * | 15.9 |
| N profile 1 | PG-HF | 0.82 | * | 0.88 | 0.97 | * | 0.84 | 10.7 | * | 15.2 |
| N profile 1 | PG-LF | 0.81 | * | 0.88 | 0.97 | * | 0.84 | 10.9 | * | 15.3 |
| N profile 2 | SG-HF | 0.76 | * | 0.87 | 0.96 |  | 0.91 | 13.1 | * | 9.8 |
| N profile 2 | SG-LF | 0.76 | * | 0.86 | 0.95 |  | 0.96 | 13.1 | * | 9.9 |
| N profile 2 | PG-HF | 0.77 | * | 0.87 | 1.02 | * | 0.9 | 12.4 | * | 9.3 |
| N profile 2 | PG-LF | 0.77 | * | 0.86 | 0.98 |  | 0.91 | 12.2 | * | 9.7 |
| E profile 1 | SG-HF | 0.76 | * | 0.88 | 0.91 | * | 0.86 | 10.5 | * | 12.7 |
| E profile 1 | SG-LF | 0.78 | * | 0.88 | 0.98 |  | 0.88 | 9.7 | * | 12.3 |
| E profile 1 | PG-HF | 0.77 | * | 0.89 | 1.08 | * | 0.91 | 10.8 | * | 13.3 |
| E profile 1 | PG-LF | 0.79 | * | 0.9 | 1.11 | * | 0.93 | 10.2 | * | 13.0 |
| E profile 2 | SG-HF | 0.78 | * | 0.86 | 1.23 | * | 0.92 | 10.4 | * | 8.6 |
| E profile 2 | SG-LF | 0.78 | * | 0.86 | 1.13 | * | 0.85 | 11.5 | * | 9.1 |
| E profile 2 | PG-HF | 0.8 | * | 0.88 | 1.06 |  | 1.02 | 10.3 | * | 8.3 |
| E profile 2 | PG-LF | 0.8 | * | 0.89 | 1.25 | * | 1.03 | 11.1 | * | 9.6 |

**Figure S3:**
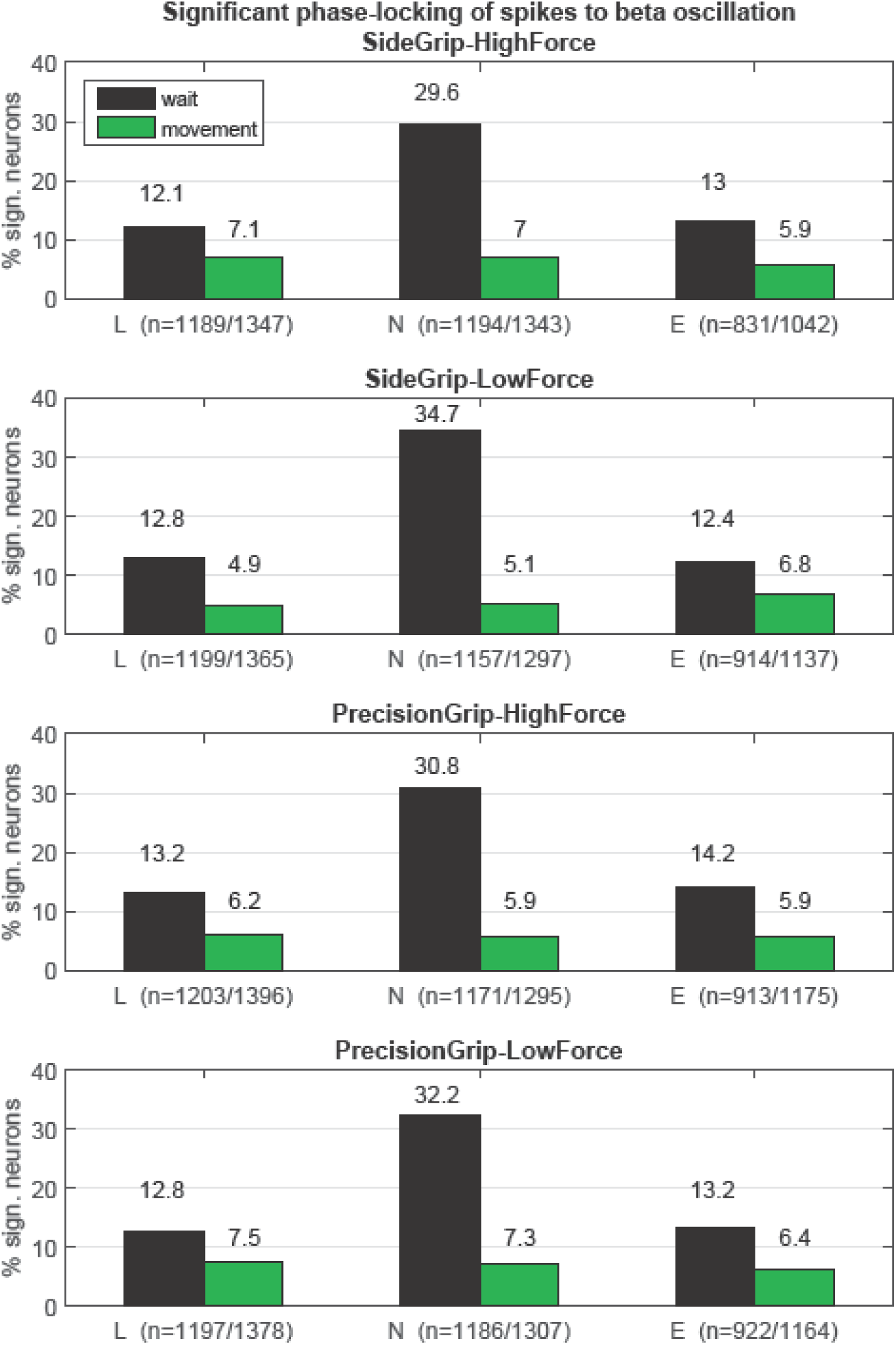
Percentages of neurons whose spike times are significantly (p<0.05) phase-locked to beta LFP oscillations in each epoch, trial type and monkey (L, N and E). Below each bar plot, the number of neurons analyzed during *wait/movement* are indicated in parenthesis.

**Table S3:** **Test of renewal prediction (see Fig. 7).** This table shows the percentages of neurons with a higher CV2^2^ than FF during *wait* and during *movement*, for all trial types and all monkeys. This percentage is very low during *wait*, but approaches the 50% during *movement*. Furthermore, the median of the individual ratios FF/CV22 obtained for each neuron are shown. For renewal processes this ratio should be 1. The value of 1 is roughly obtained during *movement*, especially in monkey L, but not during *wait*, where the FF was by far larger than CV22. This table shows that the results during all trial types were very similar.

| monkey | trial type | wait (%) | wait (FF/CV2<sup>2</sup>) | movement (%) | mvt (FF/CV2<sup>2</sup>) |
| --- | --- | --- | --- | --- | --- |
| **L** | SG-HF | 20 | 1.43 | 55 | 0.96 |
|  | SG-LF | 20 | 1.46 | 52 | 0.97 |
|  | PG-HF | 23 | 1.36 | 59 | 0.93 |
|  | PG-LF | 22 | 1.38 | 54 | 0.96 |
| **N** | SG-HF | 17 | 1.6 | 40 | 1.11 |
|  | SG-LF | 19 | 1.6 | 39 | 1.14 |
|  | PG-HF | 15 | 1.65 | 36 | 1.15 |
|  | PG-LF | 15 | 1.69 | 36 | 1.19 |
| **E** | SG-HF | 13 | 1.95 | 32 | 1.22 |
|  | SG-LF | 13 | 1.92 | 34 | 1.21 |
|  | PG-HF | 10 | 1.83 | 31 | 1.24 |
|  | PG-LF | 8.2 | 2.05 | 35 | 1.22 |

